# Functional Characterization of Myeloid Neoplasm-associated DDX41 Variants Reveals Pathogenic Interaction with Acquired Hotspot Mutation

**DOI:** 10.64898/2026.05.27.727893

**Authors:** Julianna S. Fisher, Emily Stepanchick, Andrew Wilson, Junichiro Kida, Mike Adam, Mariela V. Perez Otero, Talha Badar, Yael Kusne, Alejandro Ferrer, Mrinal M. Patnaik, Timothy M. Chlon

## Abstract

Germline variants in *DDX41* are the most frequent genetic predisposition to adult hematologic malignancies. The most common variants are truncating, implicating loss of function in the pathogenesis. However, non-truncating variants account for 30-40% of cases, and their impact on essential *DDX41* functions remains unknown. We utilized a genetic complementation assay to assess the functionality of 10 recurrent germline non-truncating variants of DDX41. All variants restored viability to *Ddx41*-deficient hematopoietic progenitor cells at exogenous expression levels. In contrast, the hotspot mutant p.R525H, which is somatically acquired at disease onset in >50% of patients, failed to restore viability. CRISPR-based modeling in cell lines and mice revealed heterogeneity: some variants were non-functional at endogenous expression levels whereas others maintained complete functionality, supporting normal cell proliferation and even lifelong hematopoiesis in a homozygous setting. Notably, co-expression of p.R525H with some variants caused impaired hematopoietic progenitor cell viability, indicating a dominant-negative effect of p.R525H. In contrast, other variants, all classified as variants of unknown significance, were unaffected by the presence of p.R525H. A screen of 100 disease-associated variants confirmed that many non-truncating germline variants are susceptible to p.R525H-mediated dominant-negative effects, whereas wild-type DDX41 is not. These findings indicate that *DDX41* variant curation is complicated by variable effects on functionality and variant-specific interactions with somatically-acquired *DDX41* mutations. The dominant-negative effect of p.R525H provides a mechanistic basis for the conclusion of recent patient cohort analyses that co-occurrence with a somatic hotspot mutation is a reliable indicator of DDX41-driven disease in carriers of non-truncating variants.

## Introduction

Heterozygous germline variants of *DDX41* are the most prevalent cause of genetic predisposition to adult myeloid malignancies, accounting for 2-4% of overall incidence of myelodysplastic syndrome (MDS) or acute myeloid leukemia (AML).^1-4^ Recent population studies indicate that as many as 1 in 430 individuals carry a Pathogenic/Likely Pathogenic (P/LP) *DDX41* variant.^5^ The average age of disease onset in these individuals is similar to the general population (approximately 70 years); however, their disease has several defining features including hypocellular marrow, the presence of a somatic (second-hit) *DDX41* mutation, and a paucity of signaling driver mutations like RAS and FLT3.^1-3,5,6^

*DDX41* codes for an essential RNA helicase that has many ascribed functions, including roles in pre-mRNA splicing, R-loop regulation, ribosome biogenesis, and as a cytosolic innate immune sensor.^7-13^ At least one of these functions is essential for development since loss of *DDX41* causes embryonic lethality and cell viability defects in proliferative cell types, including hematopoietic progenitor cells (HPCs).^7,14,15^ Due to the varied nature of its functions, the precise mechanism by which *DDX41* mutations promote the onset of MDS/AML pathogenesis remains in question, causing challenges for the assignment of pathogenicity to variants.^16-19^

Truncating or start-loss *DDX41* germline variants, which cause loss of protein expression, account for the majority of DDX41-mutated MDS/AML cases, indicating that heterozygous loss-of-function mutations cause MDS/AML predisposition.^3^ However, non-truncating germline variants account for 30-40% of cases and affect amino acids throughout the length of the protein, with more than 100 different variants being reported in disease cases.^1-3^ These variants have an undetermined impact on protein function, and thus, their role in disease compared to truncating variants is poorly understood. Previous reports have shown that both truncating and non-truncating P/LP variants (defined by American College of Medical Genetics scoring) are associated with *DDX41*-specific disease features, including co-mutation of the contralateral *DDX41* allele.^2,3^ In contrast, variants of unknown significance (VUS) are less associated with these features.^2,3^ The acquired co-mutations of *DDX41*, of which p.R525H accounts for 50-70%, are always non-truncating.^2,3^ Our previous studies indicate that p.R525H lacks the essential DDX41 function and cannot support HPC proliferation.^7^ However, the frequency with which this mutation occurs and the absence of acquired truncating mutations suggest that p.R525H has a specific effect on DDX41 function that is distinct from complete loss and promotes pathogenesis specifically in the context of existing *DDX41* mutation.

To determine the role of germline non-truncating variants in DDX41-mutated malignancy, we characterized the functional effect of 7 P/LP and 3 VUS on DDX41 function through genetic complementation assays and CRISPR gene editing in cell lines and mice. Through combined expression with the p.R525H hotspot mutant, we uncovered dominant-negative effects of p.R525H that are variant-specific, potentially explaining the role of otherwise functional variants in predisposition to the DDX41-mediated pathogenesis.

## Methods

### Cell Lines

Mouse MA9 cells were derived from Ddx41^flox^ Rosa26-CreERT2 lineage-negative bone marrow cells as previously described^7^. MA9 cell lines were cultured in IMDM with 10% FBS, 1% Penicillin-streptomycin, and 10ng/mL of hIL-6, mSCF, and mIL-3 with passaging every 2-3 days. To induce Cre activity, cells were treated for 24 hours with 1μM 4-hydroxytamoxifen (Sigma, H-7904) or ethanol (1:1000) with a media change to remove the tamoxifen. Platinum-E cell lines used for virus production were cultured in DMEM with 10% FBS, 1% Penicillin-streptomycin, 1μg/mL puromycin, and 10μg/mL blasticidin. MOLM13 cell lines were obtained from AddexBio Cat# C0003003 and cultured in RPMI 1640 with 10% FBS and 1% Penicillin-streptomycin with passaging every 2-3 days.

### Mice

Ddx41^flox^ Rosa26-CreERT2 were described previously^7,14^. Missense mutant knock-in mice (p.G173R, p.Y259C, and p.I396T) were generated by the Transgenic Animal and Genome Editing Core at Cincinnati Children’s Hospital Medical Center using CRISPR. Ribonucleoprotein (RNP) complex was formed by mixing 60ng/μL sgRNA with 80ng/μL Cas9 Protein (IDT) in Opti-MEM (Thermofisher) and incubated at 37C for 15 minutes. ssDNA donor oligos (Ultramer from IDT) with asymmetrical homologous arm design and the intended mutations were added to the RNP complex at 220ng/μL each. Zygotes from superovulated female mice on C57BL/6J background were electroporated with 7.5μL RNP/donor oligo mix on ice using a Genome Editor electroporator (BEX; 30V, 1ms width, and 5 pulses with 1s interval). Two minutes after electroporation zygotes were moved into 500μL cold M2 medium (Sigma), warmed to room temperature, and transferred into the oviductal ampulla of pseudopregnant CD-1 females. Pups were born and genotyped by PCR and Sanger sequencing.

Animals were bred and housed in the Association for Assessment and Accreditation of Laboratory Animal Care-accredited animal facility of Cincinnati Children’s Hospital Medical Center. All mouse work was conducted under an IACUC-approved protocol (2022-0055). All mouse strains were kept on C57BL/6N background. When mice were moribund (pale, extreme lethargy, panting, etc), they were sacrificed by CO2 asphyxiation, and bones and spleen were harvested for analysis. BM was harvested by crushing leg bones with a mortar and pestel. RBC were lysed in 1X Lysing Buffer (BD Biosciences). For longitudinal studies, mice were bled monthly by submandibular puncture to obtain 50-100µl of blood.

### Genotyping

DNA was isolated from mouse tails with 50mM NaOH for 1hr at 95°C. 100mM Tris HCL was added and samples were vortexed until tails dissolved. Tail debris was pelleted by centrifuging samples for 2min at 15,000xg. For Ddx41^flox^ Rosa26-CreERT2 mice, 1-2uL of DNA was genotyped for DDX41 (CSD-Ddx41-ttR2, CSD-Ddx41-F2), Rosa26-CreERT2 (CreER, Rosa26-R, Rosa26-F2) using GeneAmp Fast PCR master mix 2x (ThermoFisher).

Missenese mutant mice (p.G173R, p. Y259C, and p.I396T) were genotyped using mutation-specific restriction digest. PCR products were first amplified and purified using the QIAquick PCR Purification Kit (Qiagen, 28106), followed by restriction digest. The G173R amplicon was digested with BsbrI (NEB, R0102S), Y259C with DpnII (NEB, R0543T), and I396T with HpyCH4IV (NEB, R0619L). Digested products were resolved on a 3% agarose gel at 130V for 1 hour and visualized using the BioRad ChemiDoc.

### CRISPR Missense Mutant Cell Lines

To generate DDX41 missense mutations at the endogenous locus in MOLM13 cells, sgRNA:Cas9 complexes were prepared using 1.5μg Alt-R S.p.Cas9 (IDT, 1081058) with 1μg sgRNA (Synthego) and incubated at room temperature for 20-30 minutes. Following incubation, 60-100μM donor Oligo (IDT) was added to sgRNA:Cas9 complex, which was then transfected into ∼200,000 cells using the NEON transfection system (Thermo, settings: 1400V, 20ms, 1 pulse). Media was changed 24 hours post-transfection, and single cells clones were plated in a 96-well u-bottom plate (Neta Scientific, 351177) and grown in a humidity chamber. To screen single cells clones, genomic DNA was isolated, PCR-amplified and purified by gel electrophoresis, followed by gel extraction (Zymo, C1004-50) prior to Sanger sequencing. RNA was isolated using the Quick RNA MiniPrep Kit (Zymo, R1055), and cDNA was synthesized using the High-capacity cDNA Reverse Transcription (Life Tech,4368814). The presence of the intended missense mutations was further confirmed by Sanger sequencing.

### Plasmids and Transduction

Generation of MSCV-mCherry-HA-DDX41 and MSCV-mCherry-HA-DDX41 R525H from MSCV-IRES-mCherry vector (Addgene, #52114) was previously described. To generate plasmids containing DDX41 variants (p.K9E, Y33H, D140fs, G173R, M155H, R164W, R219H, P258L, and I396T), site-directed mutagenesis of MSCV-mCherry-HA-DDX41 plasmid was performed using the QuikChange II XL Site-Directed Mutagenesis Kit (Agilent, 200521) and mutation-specific primers. DDX41 variants Q52fs and K331del mutant plasmids were purchased from Twist Biosciences on MSCV-mCherry-HA-DDX41 backbone. For co-expression experiments, mCherry was replaced with BFP by cloning into MSCV-IRES-mCherry vector at NcoI and Notl sites. HA-DDX41 and HA-DDX41 R525H were then subcloned into MSCV-IRES-BFP vector at EcoRI and BamHI sites. All plasmid sequences were confirmed by Sanger sequencing.

Retroviral supernatants were made by transfecting PlatE cells using TransIT-LT1 (Mirus) transfection reagent with 24 μg retroviral plasmid and harvesting the culture medium 48 hours post-transfection. MA9 Cells were transduced with viral supernatant containing 0.8 μg/ml polybrene for 24 hours, and expression of BFP or mCherry was assessed by flow cytometry. For Lin^-^ cells, viral media was spun on retronectin coated plates at 800xg for 2 hours at 32°C. Following centrifugation, viral supernatant was removed and Lin^-^ cells in fresh media were added to the plates and incubated at 37°C overnight. The following day, cells were collected and media was replaced. Seventy-two hours after transduction, mCherry or BFP positivity was assessed via flow cytometry, and cells were sorted for downstream analysis.

### DDX41 Variant Library

A DDX41 variant library was generated containing germline and somatic DDX41 variants observed in patients^1-3^. Each plasmid contained a unique barcode after the stop codon. Retroviral supernatants were produced by transfecting Platinum-E cells using TransIT-LT1 (Mirus) transfection reagent with 24 μg retroviral plasmid and harvesting the culture medium 48 hours post-transfection. DDX41 variant library viral media was co-transfected with DDX41 R525H or empty vector control on retronectin coated plates were spun in viral media at 800xg for 2 hours at 32°C. Following centrifugation, viral supernatant was removed and MA9 cells in fresh media were added to the plates and incubated at 37°C overnight. The next day, cells were removed from plates and transferred to fresh media. Seventy-two hours after transduction cells were sorted for mCherry and BFP positivity. Sorted cells were allowed to recover for 48 hours prior to collection of the day 0 timepoint and treatment with 1μM 4-hydroxytamoxifen for 24 hours. Genomic DNA was collected on Day 0 and Day 14 after tamoxifen treatment and purified using Quick-DNA Miniprep Plus Kit (Zymo, D4068). A PCR product containing the barcode was generated using ExTaq Polymerase (Clontech) and primers that included the P7 and P5 sequences for Illumina sequencing. Products were sequenced on an Illumina Novaseq, and the abundance of each barcode relative to wild-type control plasmid was determined. Short-read sequencing data were imported into R from fastqs using the ShortRead^20^ and Biostrings^21^ packages. Reads were screened for the presence of a target motif consisting of defined left, CTCCATGGACTTCTGA, and right, AGCGGTGTTTCG, flanking sequences separated by a variable 5-nt region. Approximate matching was performed with vmatchPattern, allowing up to one mismatch across the full motif and disallowing indels. For each matched read, the coordinates corresponding to the 5-nt variable region were extracted using extractAt and tabulated to obtain counts of unique variants.

### Proliferation Assay

MA9 cells were treated with 1μM 4-hydroxytamoxifen or ethanol (1:1000) for 24 hours. Following treatment, the media was replaced, and cells were counted using trypan blue on a Countess Automated Cell Counter (ThermoFisher, C10228) and plated at 200,000/mL. Cells were counted daily for 7 days and passaged to maintain a density of ∼500,000 cells/mL. MOLM13 cells were plated at 300,000 cells/mL and similarly counted in trypan blue on the Countess for 7 days, with passaging to maintain ∼500,000 cells/mL. At 48 hours, cells were collected for RNA isolation, and at 72 hours, cells were collected for protein extraction and Annexin V analysis.

### Quantitative PCR

RNA was collected from MA9FF cells 48 hours after treatment with 1μM 4-hydroxytamoxifen or ethanol (1:1000) and MOLM13 cells were collected 48 hours after plating cells for proliferation assays, using the Quick RNA MiniPrep Kit (Zymo, R1055). A total of 1μg RNA was used for cDNA synthesis with the High capacity cDNA Reverse Transcription (Life Tech,4368814). cDNA was diluted 1:4, and samples were run in triplicate on a 96-well Fast Thermal Cycling plate (Life Tech, 4346907) using Powerup SYBR Green Master Mix (ThermoFisher, A25777).

### Immunoblotting

Whole cell lysates were made with RIPA buffer (20mM Tris HCL pH7.4, 37mM NaCL, 2mM EDTA, 1% Triton X-100, 10% Glycerol, 0.1% SDS, 0.5% NaDeoxycholate) in the presence of PMSF (10mM final), complete Mini Protease Inhibitor Cocktail (Roche), and Phosphatase Inhibitor Cocktail 2 and 3 (Sigma Aldrich). Lysates were separated by SDS-PAGE, transferred to nitrocellulose membranes, and immunoblotted. Immunoblots were performed with the following antibodies: DDX41 (Novus, H00051428), Vinculin (E1E9V) XP® Rabbit mAb (Cell Signaling, 13901), and GAPDH (D16H11) XP Rabbit mAb (Cell Signaling, 5174).

### Colony assay

Lineage negative cells were isolated from Ddx41^flox^ Rosa26-CreERT2 with Miltenyi Mouse Lineage Cell Depletion Kit (130-110-470). Lin^-^ cells were transduced on retronectin-coated plates and sorted for mCherry and/or BFP positivity using a Sony SH800S Cell Sorter. For single mutant experiments, cells were plated at 1000 Lin^-^ cells per well in triplicate in MethoCult (STEMCELL) M3434 (myeloid) and 1μM 4-hydroxytamoxifen was added to the MethoCult. Colonies were counted after 12 days using Stemvision (StemCell Technologies). For co-expression of P/LP and p.R525H variant experiments 4000 Lin^-^ cells per well were plated in triplicate and treatments were conducted the same as single mutant experiments.

### Flow Cytometry

MA9FF cells were treated with 1μM 4-hydroxytamoxifen or ethanol (1:1000) for 72 hours and stained for Annexin V (1:100) (eBiosciences, 88-8007-74). MOLM13 cells were collected for Annexin V staining 72 hours after plating for proliferation assays. All flow cytometry analyses were performed on LSRII (BD Biosiences), and data were analyzed using FlowJo Software.

## Statistics

Pairwise-comparisons were calculated by T-test, and multiple comparisons were calculated by ANOVA followed by Dunnett’s test for individual comparisons. Significance was determined on a 95% confidence interval. Contingency analysis on mouse genotype outcomes was conducted by Chi-square test. Significant differences are noted by * (<0.05), ** (<0.01), **** (<0.0001).

## Results

### Germline DDX41 missense variants rescue the growth of *Ddx41*-deficient HSPCs

To determine the functional effect of germline non-truncating *DDX41* variants on protein function, we selected seven recurrent non-truncating P/LP variants (R164W, G173R, R219H, P258L, Y259C, K331del, and I396T) and three recurrent VUS (K9E, Y33H, M155H) for characterization (Figure 1A). Predictive structural modeling of these variants indicates that most P/LP variants result in minor, localized structural differences within the core of the DEAD domain, without disrupting global protein structure.^22^ Notably, they are situated away from the ATP binding pocket, which is formed by the interface of the DEAD and Helicase domains and contains sites of common acquired mutations including R525H and others (e.g. T227M, G530D).^22^ In contrast, all three VUS and p.I396T are found on the exposed protein surface and do not result in alterations to either global or local protein structure (Figure 1B; Figure S1A).^22^ Since DDX41 loss of function is lethal to proliferative cells, including HPCs, the functionality of variant protein can be assessed by the ability to rescue the growth of *Ddx41*-deficient HPC.^7^ We utilized Lin-BM cells isolated from *Ddx41*^flox/flox^ Rosa26-CreERT2 mice, enabling tamoxifen-inducible loss of Ddx41 expression, which is lethal to the cells.^7^ We retrovirally transduced *Ddx41*^flox/flox^ Lin-BM with the 7 P/LP variants, 3 VUS, 2 recurrent truncating variants (p.Q52fs and p.D140fs), the recurrent somatic mutation p.R525H, and wild-type DDX41, and then plated the cells in methylcellulose (Figures 1C and 1D). Surprisingly, we observed that all non-truncating germline variants (P/LP and VUS) rescued the colony forming potential of *Ddx41*-deficient Lin-cells, with only one variant (R164W) showing a significant difference to wild-type (Figure 1E). In contrast, truncating *DDX41* variants and the somatic p.R525H mutant failed to rescue colony forming potential, indicating loss of function. Concurrently, in an immortalized *Ddx41*^flox/flox^;RosaCreERT2 cell line, which also undergoes cell death upon tamoxifen treatment, we observed that exogenous expression of non-truncating germline variants were sufficient to rescue the proliferation and viability of *Ddx41*-deficient cells (Figures 1F and 1G; Figures S1B and S1C). We again found that R164W had a significant reduction in cell growth compared to wild-type although viable cells were present. In contrast, both truncating and p.R525H variants were completely incapable of rescuing the proliferation and viability of these cells. Previously, we described that loss of Ddx41 causes defects in splicing at introns that contain snoRNAs, resulting in aberrant intron retention. To determine the ability of P/LP variants and VUS to regulate splicing at snoRNA-containing introns, we performed targeted qPCR for the retention of *Snora7a* within the host gene *Rpl32*. We found that all germline non-truncating variants were equally as capable as WT to properly regulate splicing at the Rpl32-Snora7a locus whereas p.R525H and truncating DDX41 variants exhibited increased intron retention indicative of the splicing defect characteristic of DDX41 loss of function (Figure 1H). These studies demonstrate that non-truncating P/LP and VUS *DDX41* variants, but not recurrent somatic mutation p.R525H and truncating DDX41 variants, are able to maintain essential DDX41 functions in HSPCs.

**Figure 1.**
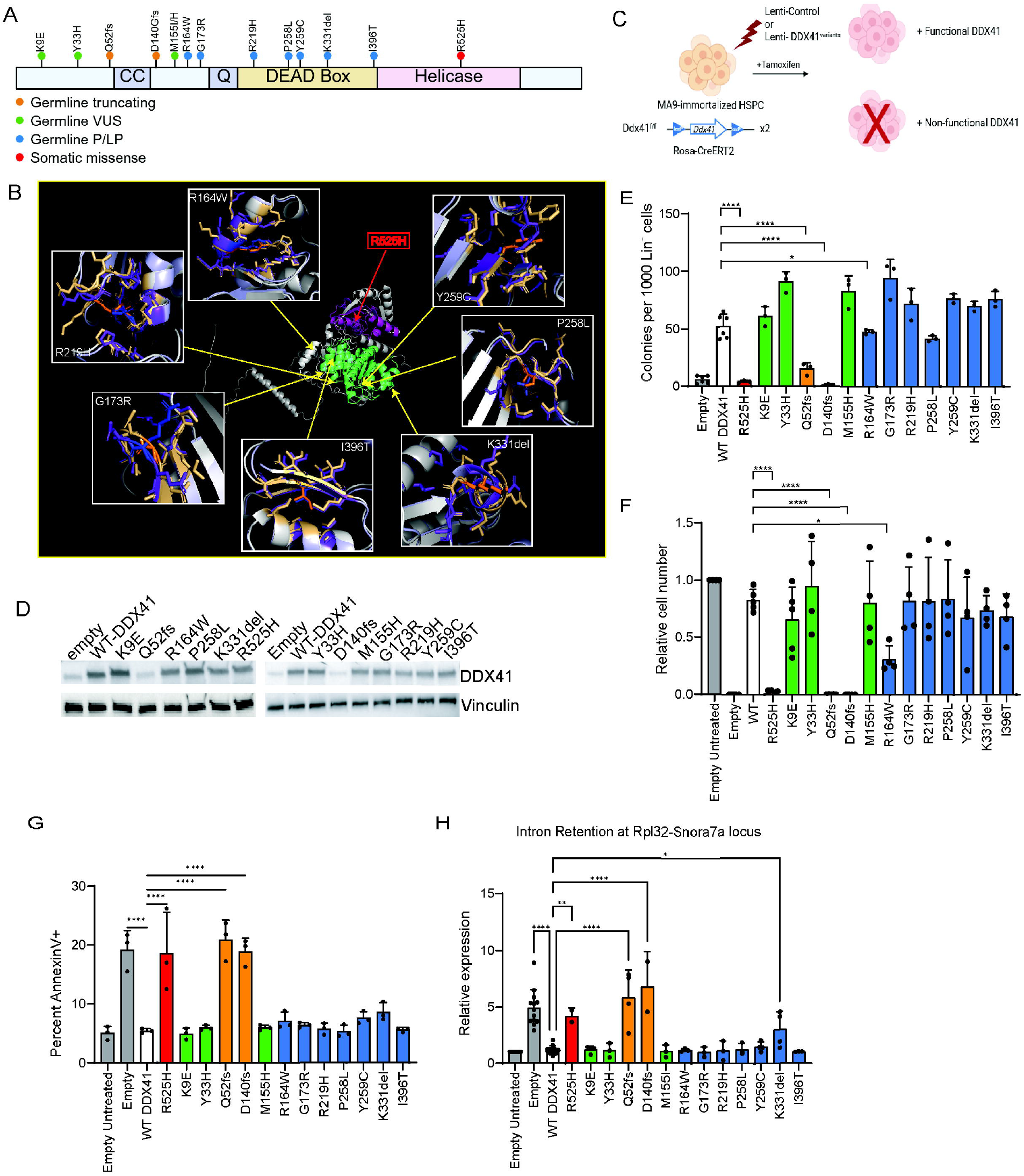
Exogenous expression of disease-associated missense mutants of DDX41 rescues growth of *Ddx41*-deficient hematopoietic cells. (A) Non-truncating P/LP and VUS germline variants selected for characterization. (B) Predictive modeling with AlphaFold 3 of non-truncating P/LP and wild-type DDX41 variants generated with PyMOL. The center is a ribbon diagram of DDX41 with the DEAD domain in green and the Helicase domain in purple. The location of P/LP variant residues is indicated by arrows. Each inset includes a ribbon diagram showing the wild-type protein in gold and the variant protein in purple. The affected wild-type amino acid is shown in orange and the variant amino acid in blue. (C) Schematic of *Ddx41*-deficient Lin^neg^ rescue experiment. (D) Immunoblot for DDX41 expression in Lin^neg^ cells transduced with lentivirus expressing the indicated DDX41 variant. (E) Myeloid colony assay on cells from B. (F) Proliferation of an immortalized *Ddx41*^flox/flox;^ RosaCreERT2 cell line expressing indicated DDX41 variant 7 days after tamoxifen treatment. (G) Annexin V staining of immortalized *Ddx41*^flox/flox^ cells 72 hours after tamoxifen treatment. (H) Intron retention at the Rpl32-Snora7a locus in immortalized *Ddx41*^flox/flox^ cells 48 hours after tamoxifen treatment.

### Homozygosity of germline non-truncating P/LP variants and VUS at endogenous expression levels reveals a hypomorphic phenotype

Given that exogenous expression of DDX41 P/LP and VUS variants were sufficient to rescue the growth of *Ddx41*-deficient cells, we sought to determine the ability of these variants to maintain DDX41 function at endogenous expression levels. To do so, we utilized CRISPR homology-directed repair (HDR) to modify the *DDX41* alleles in MOLM13 cells, a human AML cell line, which we confirmed to be diploid for *DDX41* (Figure 2A, Figure S2A). To isolate clones with homozygous mutations, CRISPR-edited cells were plated as single cells, and the resulting clones were screened by Sanger sequencing (Figure 2A; Figure S2A). We produced viable, proliferating cell lines with homozygous mutations for all 3 VUS and 4 of 7 P/LP variants (Figure 2B; Figure S2A). For R164W, P258L and K331del, we were unable to produce homozygous mutant cell lines despite screening hundreds of clones and finding several with heterozygosity. By western blot, we confirmed that the homozygous mutant clones expressed DDX41 protein and observed that K9E and Y33H-mutated clones had increased expression of DDX41 while the other clones had similar expression as parental cells (Figure 2C). We monitored the cell number in the culture over 7 days and found that R219H and Y259C-mutated clones had reduced proliferation while clones with other P/LP variants or VUS had similar proliferation to wild-type cells (Figure 2D; Figure S2B). Since 3 of 7 P/LP variants could not be generated as homozygous modifications and 2 more caused reduced cell proliferation, we concluded that some germline P/LP variants cause hypomorphic effects on DDX41 function. G173R and I396T were unique in that they supported the proliferation and viability of MOLM13 cells equally to wild-type.

**Figure 2.**
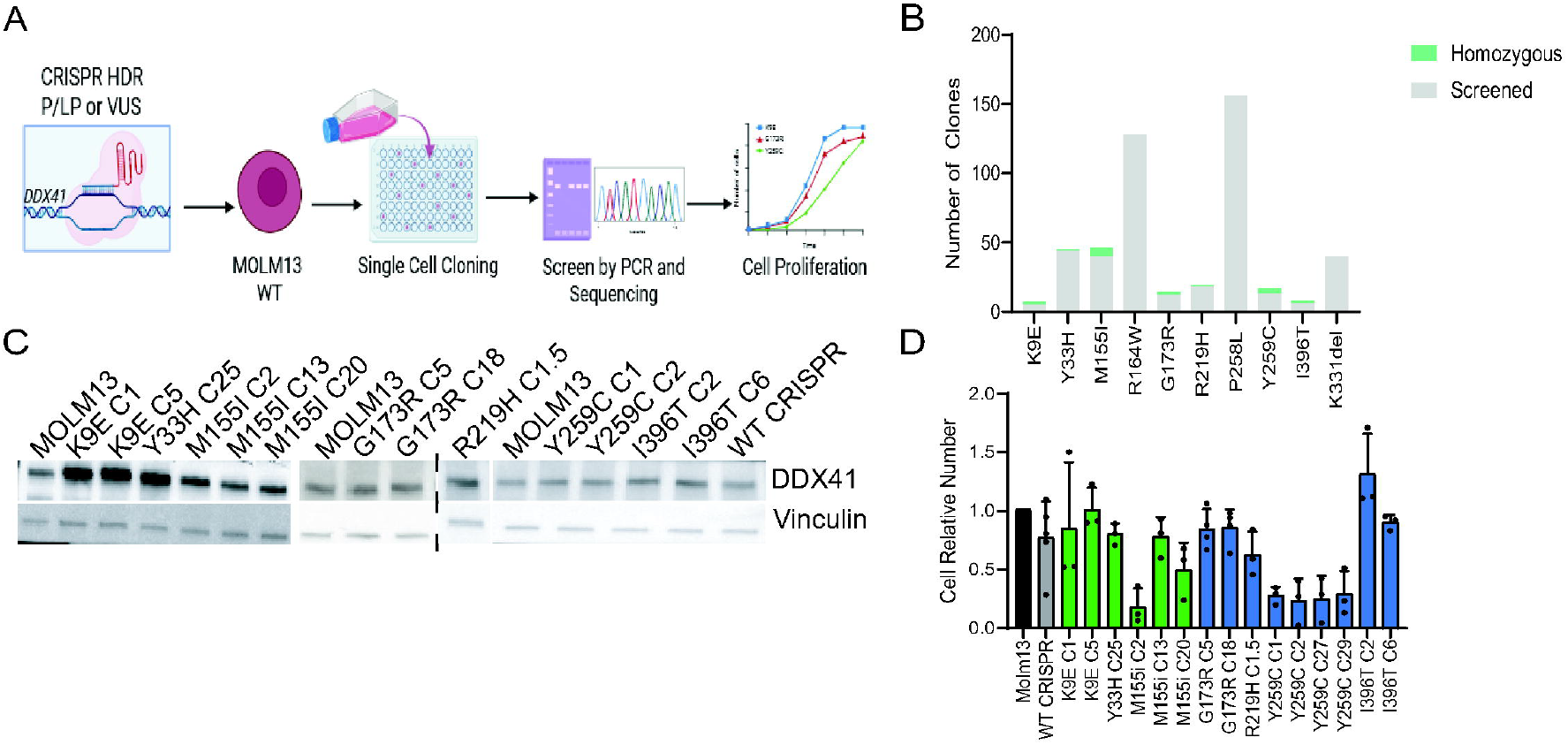
Homozygosity of disease-associated mutations in the endogenous *DDX41* alleles reveals hypomorphic effect on protein function. (A) Schematic of CRISPR targeting of wt-MOLM13 cells in endogenous *DDX41* loci. (B) Number of homozygous lines generated from colonies screened in CRISPR targeting of MOLM13 cells. (C) Immunoblot for DDX41 expression in MOLM13 cells with homozygosity for the indicated mutation. (D) Proliferation of homozygous *DDX41*-mutant cell lines normalized to wt-MOLM13 cells.

### Generation of p. G173R, p. Y259C, and p.I396T *Ddx41* alleles in mice reveals variable effects on viability

To further characterize P/LP variants with varying degrees of functional deficit, we generated C57Bl/6 mouse lines harboring p.G173R, p. Y259C, and p.I396T mutations in the endogenous *Ddx41* alleles (Figure 3A). Heterozygous p.G173R, p.Y259C, and p.I396T mice exhibited complete blood cell counts similar to wild-type littermates up to one year of age, despite a modest reduction in Ddx41 protein expression (Figures 3B and 3C). However, homozygous p.G173R and p.Y259C were not obtained across many litters from multiple breeding pairs. Chi-square analysis confirmed the observed lack of homozygous pups compared to expected, indicating that homozygosity of these P/LP variants causes lethality during embryonic or fetal development. In contrast, p. I396T homozygous mice were viable and born at Mendelian ratios. Although homozygous p.I396T mice showed a marked reduction in DDX41 protein expression compared to both wild-type and heterozygous littermates, they remained healthy and maintained normal blood counts up to at least 15 months of age (Figures 3B and 3C).

**Figure 3.**
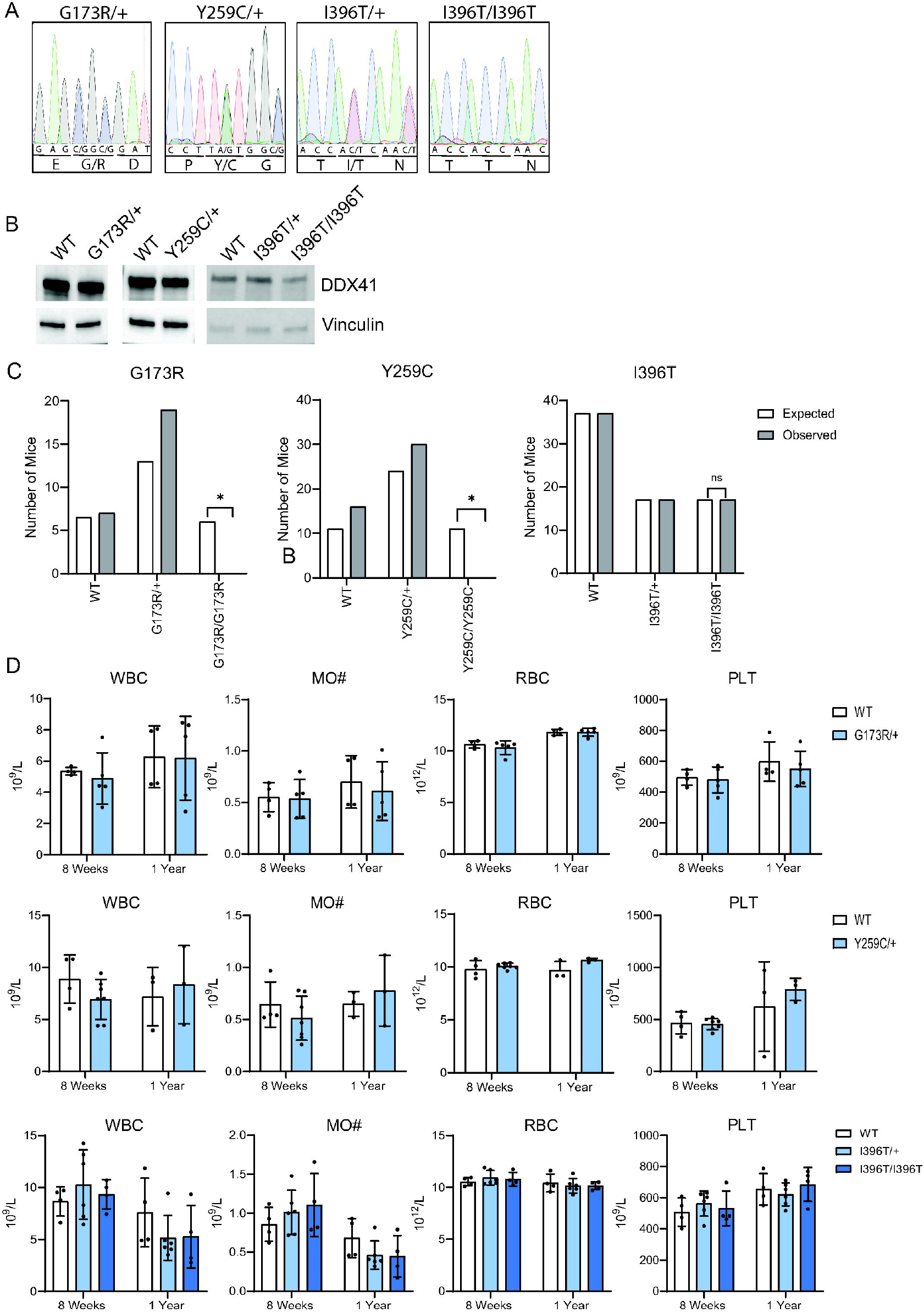
Heterozygosity of pathogenic/likely pathogenic Ddx41 variants (p.G173R, p.Y259C, and p.I396T) is tolerated and results in phenotypically normal mice. (A) Sanger sequencing of p.Y259C, p.G173R, and p.I396T heterozygous and p.I396T homozygous mice. (B) Immunoblot for DDX41 expression in Lin^neg^ cells shows decreased DDX41 expression. (C) Comparison of observed versus expected numbers of homozygous pups of p.Y259C, p. G173R, and p.I396T mice. (D) Complete blood count (CBC) on heterozygous p.Y259C, p.G173R, and p.I396T and homozygous p.I396T mice at 8 weeks and 1 year old.

### The somatic hotspot mutant p.R525H has dominant negative effects on non-truncating P/LP DDX41 variants

Given the prevalence of somatic p.R525H mutations in patients with germline non-truncating P/LP *DDX41* variants, we sought to model the combined effect of these mutations. Using the previously described Lin-BM cells with inducible deletion of *Ddx41*, we co-expressed each non-truncating P/LP and VUS with p.R525H by viral transduction and measured the colony forming potential. We found that all P/LP variants and p.M155H had dramatically reduced colony formation when combined with p.R525H expression whereas WT DDX41, p.K9E, and p.Y33H supported normal colony formation when co-expressed with p.R525H. In the immortalized cell line system, we found that co-expression of some P/LP variants (p.G173R, p.P258L, and p. Y259C) with p.R525H resulted in reduced cell proliferation and viability compared to WT only and WT combined with p.R525H (Figures 4C and 4D). Collectively, these results indicate that expression of P/LP variants with the somatic hotspot mutation p.R525H causes HPC defects consistent with the DDX41-mutant MDS phenotype. This observation implies that the somatic variant acts in a dominant-negative manner toward many non-truncating P/LP DDX41 variants, which have varying degrees of functional deficit.

**Figure 4.**
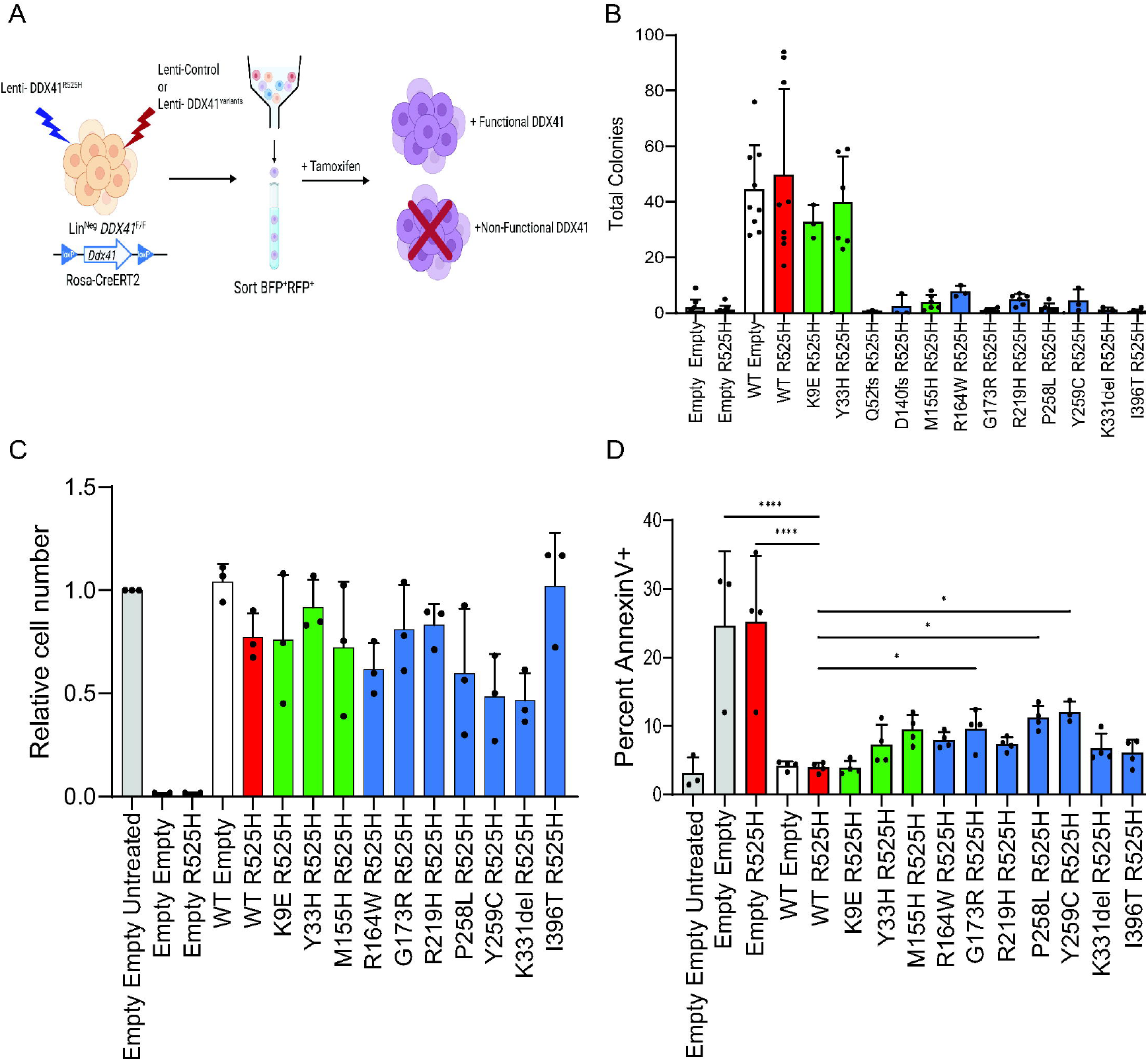
A subset of missense mutants causes progenitor cell impairment in combination with the R525H mutation. (A) Schematic of *Ddx41*-deficient Lin^neg^ rescue experiment. (B) Colony assay on Lin^neg^ cells with indicated germline variant and co-expression of p.R525H. (C) Proliferation of immortalized *Ddx41*^flox/flox^ cells expressing indicated DDX41 germline variant with co-expression of p.R525H 7 days after tamoxifen treatment. (D) Annexin V staining of immortalized *Ddx41*^flox/flox^ co-expression cells 72hours after tamoxifen treatment.

### Co-expression of p.R525H mutant with a library of disease-associated DDX41 variants reveals germline variants affected by dominant negative activity

To broadly characterize the functional effect of disease-associated DDX41 variants, we generated a library of retroviruses expressing 100 germline and somatic variants. The library is a comprehensive representation of variants identified in three recent clinical cohorts.^1-3^ Wild-type DDX41 was included as a control, and each plasmid was labeled by a distinct barcode to enable enrichment analysis by high-throughput sequencing of a vector-specific PCR product. We transduced the immortalized Lin-Ddx41^f/f^;RosaCreERT2 cell line with the library and then exposed the cultures to tamoxifen for 4 and 14 days. We determined the abundance of each variant relative to wild-type at each timepoint compared to pre-treatment. This revealed gradual depletion of the truncating variants and the somatic mutants, indicating that these cause significant loss of DDX41 function even at exogenous expression levels (Figure 5B). In contrast, the non-truncating germline variants were largely maintained at their pre-treatment levels, indicating maintained DDX41 function, recapitulating the effect observed when we expressed 10 such variants individually above. To determine the functional impact of combined expression of each germline variant with the somatic p.R525H variant, we co-transduced the cells with p.R525H and the library and then conducted the same tamoxifen-treatment regimen. In this context, we found that many germline non-truncating variants were gradually depleted over time, indicating that p.R525H has dominant-negative effects on these variants that is greater than its effect on wild-type DDX41 (Figure 5C; Figure S4). The germline truncating variants and somatic mutants were similarly depleted as in the cells without p.R525H, indicating the presence of p.R525H does not impact their functional effect since they already cause severe or complete loss-of-function (Figure 5D; Figure S4). These results broadly indicate that germline non-truncating variants do not cause severe defects in protein function but combination with p.R525H has a similar functional effect as truncating mutations combined with p.R525H, conferring proliferation defects that are a hallmark of the pathogenesis. Since overexpression masks detrimental effects of some variants (e.g. P258L and Y259C), this library screen is not likely a reliable readout of pathogenicity on its own. Nonetheless, it is demonstrative of the dominant-negative effect of acquired *DDX41* mutations that is likely critical to the pathogenic mechanism.

**Figure 5.**
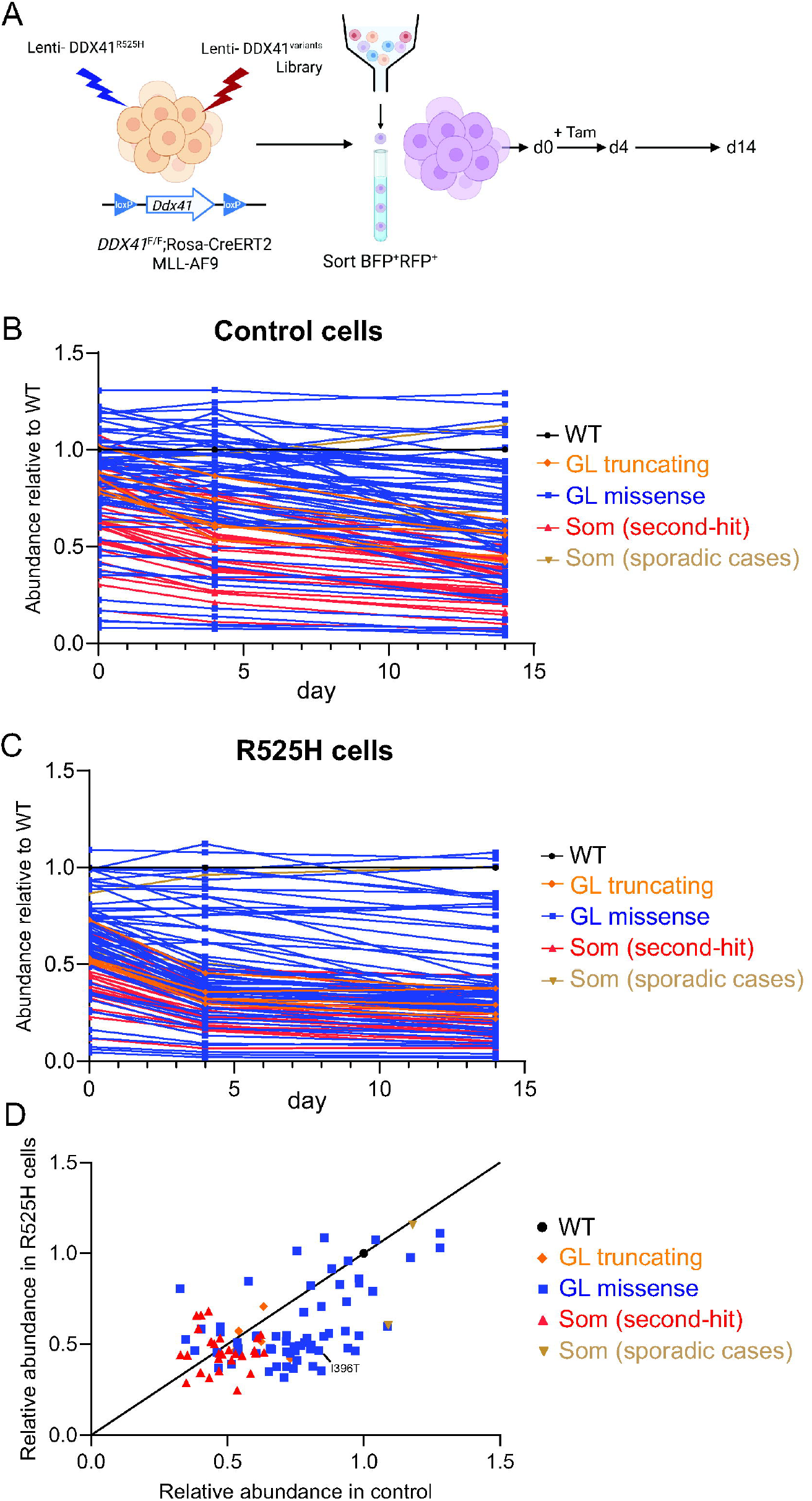
Pathogenic/likely pathogenic germline variants show reduced abundance upon co-expression with p.R525H in a *DDX41* variant library screen. (A) Schematic of *Ddx41*-deficient Lin^neg^ germline and somatic variant library screen. (B) Abundance of DDX41 germline and somatic variants relative to wild-type DDX41 in control cells. (C) Abundance of DDX41 germline and somatic variants relative to wild-type DDX41 with co-expression of p.R525H variant. (D) Relative abundance of germline and somatic DDX41 variants in control compared to p.R525H cells.

## Discussion

Our study aimed to functionally characterize a representative panel of germline non-truncating *DDX41* variants and assign pathogenicity based on loss-of-function effects. Using a genetic complementation assay in *Ddx41*-deficient cells and CRISPR-based modeling, we found that these variants affect protein function variably and incompletely. All ten non-truncating germline variants rescued the growth of *Ddx41*-deficient cells when exogenously expressed, indicating retention of partial protein function (Supplementary Table 1). This is in sharp contrast to the somatic hotspot mutant p.R525H, which is completely non-functional by the same assay. Further analysis at endogenous expression levels through CRISPR editing in both cell lines and murine models revealed clear functional deficiencies in a subset of variants. The discrepancy between these findings and the complementation assay is most likely attributable to overexpression artifacts, which can mask partial loss-of-function phenotypes, even though the proteins were no more than 2-fold overexpressed in our system (Supp 1B). Notably, several P/LP variants retained sufficient activity to sustain cell proliferation even in a homozygous-mutant context at endogenous expression levels, and one variant in particular, p.I396T, fully supported normal murine development at endogenous levels. This is especially intriguing given that p.I396T has been identified in multiple MDS/AML patients harboring a co-occurring p.R525H mutation, a hallmark of DDX41-driven malignancy^3,9^. These observations prompted us to test whether p.R525H exerts dominant-negative effects on specific germline variants. Indeed, we found that p.I396T and other P/LP variants lose functionality in the presence of p.R525H. This finding reveals a previously unrecognized dominant-negative property of the p.R525H mutant, which likely contributes to its high prevalence in DDX41-mutant malignancies.

Somatic second-hit mutations in the contralateral allele occur in more than 50% of MDS/AML patients with germline *DDX41* alterations, and over half of these involve the p.R525H hotspot. The recurrent selection of this mutation suggests a gain-of-function property that contributes to the pathogenic mechanism. Importantly, p.R525H occurs at similar frequencies in patients with either truncating or non-truncating germline mutations^3^, indicating a consistent functional role across genetic contexts. Prior studies have shown that p.R525H disrupts ATPase activity and impairs many of DDX41’s pleiotropic functions including splicing, ribosome biogenesis, and innate immune signaling^7,8,13,23^. While these findings and our complementation data are consistent with a loss-of-function effect, the absence of truncating or deletion alleles as second hits suggests that complete loss of DDX41 activity is not the effect underlying pathogenic selection of the biallelic mutant cells.

Our results indicate that p.R525H acts as a dominant negative to the non-truncating germline variants, causing HPC defects that do not occur with isolated expression of the germline variants. While this may explain the selection for the p.R525H mutation in patients with non-truncating germline variants, the equal prevalence of p.R525H in patients with truncating mutations, where a dominant negative effect is not relevant, raises further questions. One possible explanation is that p.R525H has an optimized low level of activity that is beneficial to a malignant cellular state in MDS stem cells. Testing this hypothesis will require precise, quantitative assays of DDX41 function in vivo, which remain technically challenging.

Notably, non-truncating DDX41 germline variants affect a distinct set of residues from the recurrent somatically acquired mutations, including p.R525H, p.G530D, p.P321L, p.T227M, p.A225V, which are rarely, if ever, found in the germline. These somatic mutations cluster near the ATP-binding pocket at the interface of the DEAD-box and helicase domains, whereas germline variants are situated away from this area and predominantly affect residues within the DEAD-box domain but also extend to the helicase domain and N/C-terminal regions. This difference is suggestive of distinct functional effects wherein the somatic mutations are more severe and impact ATP hydrolysis directly.

To functionally characterize a more comprehensive panel of mutants, we performed a screen of 100 disease-associated germline and somatic variants, assessing their capacity to rescue *Ddx41*-deficient cell growth. This analysis reinforced several key findings. First, germline non-truncating variants generally retain partial function sufficient to support cell growth. Second, somatic variants exhibit more severe loss-of-function phenotypes and are consequently depleted relative to wild-type controls. Third, a subset of germline variants becomes functionally compromised only in the presence of p.R525H, consistent with a dominant-negative interaction.

The molecular basis of this dominant-negative effect remains to be elucidated. Although DDX41 has not been shown to homodimerize, other DEAD-box helicases do, raising the possibility that direct protein–protein interaction underlies the observed phenotype. Alternatively, p.R525H may compete with wild-type DDX41 or variants for shared binding partners or nucleic acid substrates, thereby impairing overall functional output.

Our data represents the most comprehensive functional variant characterization to date and demonstrates that *DDX41* variant characterization should include analysis at endogenous expression levels and account for potential phenotypic modification by the p.R525H mutation. Despite the lack of a single assay to definitively demonstrate pathogenicity of individual variants, our data support the assertion of prior studies that the co-existence with a somatic hotspot mutation in MDS/AML patients is a reliable indicator of pathogenicity of a non-truncating germline variant^2,3^. This is particularly important given that DDX41-mutant MDS/AML represents a unique clinical entity that does not conform to established prognostic scoring systems^24,25^. In the absence of an acquired p.R525H mutation, even documented P/LP variants may not have direct prognostic impact. These cases should be carefully assessed based on known prognostic indicators including disease phenotype, additional co-mutations (eg. *TP53*), and other clinical variables. Beyond prognostic interpretation, our findings provide rationale for high-sensitivity screening for p.R525H mutations in known carriers of *DDX41* variants. Conversely, our study also supports confirmation of germline *DDX41* status when p.R525H or other somatic hotspot mutations are identified in patients since non-truncating germline variants may only manifest pathogenically in combination with the somatic hotspot.

Current models of DDX41-mediated pathogenesis propose that reduced DDX41 function leads to clonal dominance of biallelic-mutant HSC, resulting in HPC proliferation defects and multi-lineage cytopenias. Within this context, malignant HSC produce blasts that drive disease progression, potentially through additional genetic hits. Our study refines the molecular mechanism by demonstrating that the somatic hotspot mutation p.R525H instills a critical threshold of functional impairment through dominant-negative effects on non-truncating germline variants. Future studies of patient samples and animal models with non-truncating germline mutations are needed to shed further light on the pathogenic mechanism and to identify novel therapeutic targets.

## Supporting information

Oligo sequences

