## Supplementary material for "Functional Characterization of Myeloid Neoplasm-associated DDX41 Variants Reveals Pathogenic Interaction with Acquired Hotspot Mutation"

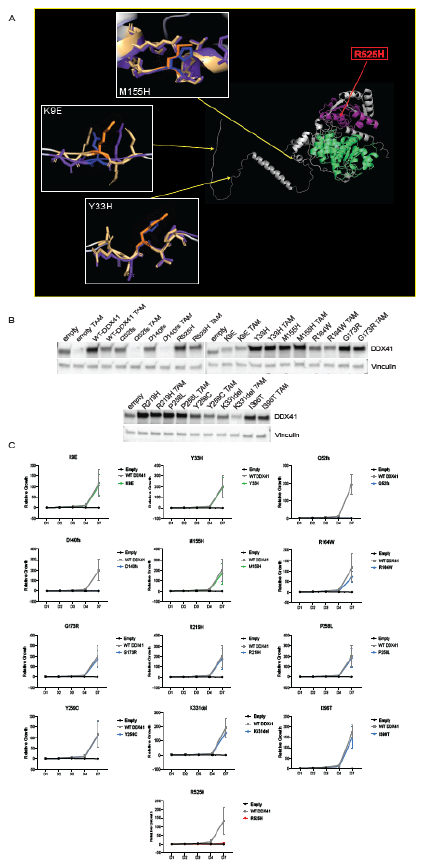

Supplementary Figure 1. (A) Predictive modeling with AlphaFold 3 of non-truncating VUS and wild-type DDX41 variants generated with PyMOL. (B) Immunoblot for DDX41 expression in immortalized *Ddx41*^flox/flox^ cells transduced with lentivirus expressing DDX41 variants. (C) Proliferation of indicated DDX41 variant compared to wild-type and empty vector expressing cells for 7 days after removal endogenous *DDX41* expression in immortalized *Ddx41*^flox/flox^ cells.

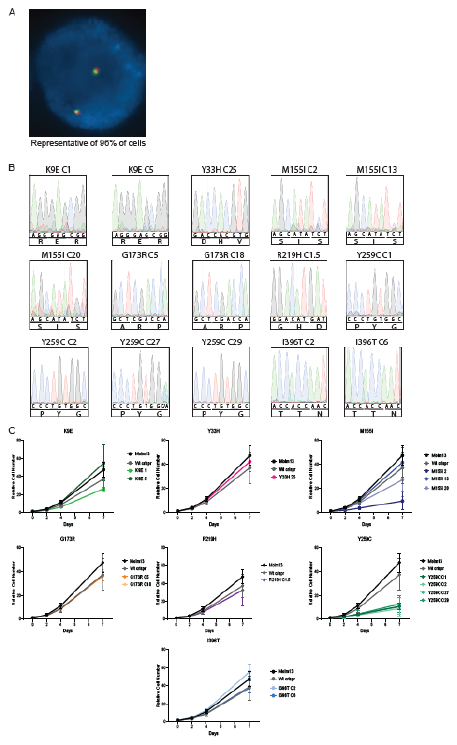

Supplementary Figure 2. (A) Fluorescence in situ hybridization (FISH) confirmation of 2 DDX41 (5q35.3) copies in parental wt-MOLM13 cells. (B) cDNA sequencing confirmation of homozygous missense mutations at endogenous *DDX41* loci. (C) Proliferation of each MOLM13 CRISPR VUS and P/LP variant over 7 days.

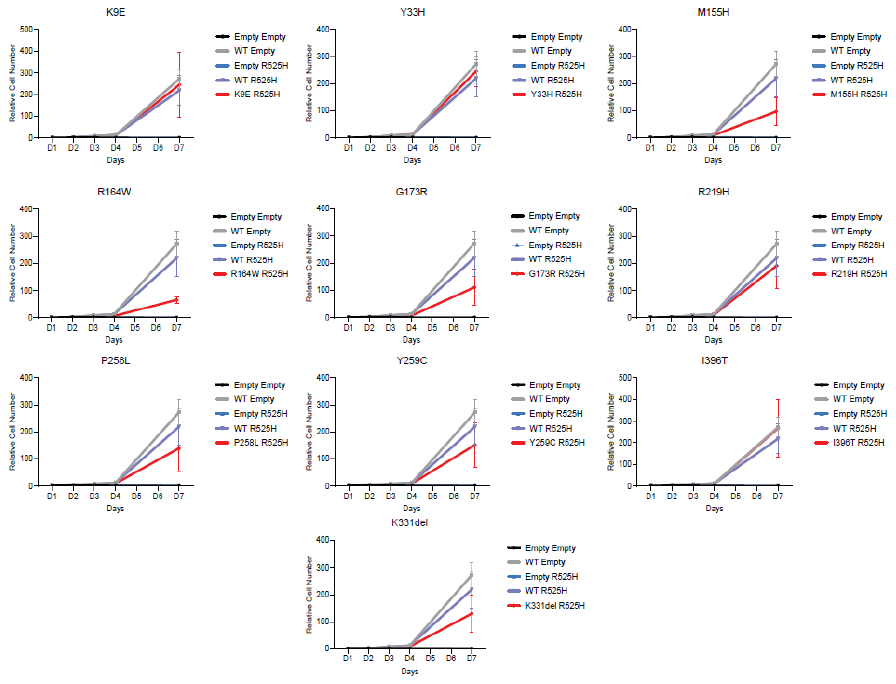
Supplementary figure 3. Proliferation of immortalized *Ddx41*^flox/flox^ cells expressing indicated DDX41 germline variant with co-expression of p.R525H over 7 days after tamoxifen treatment compared to empty and wild-type vector cells with p.R525H co-expression.

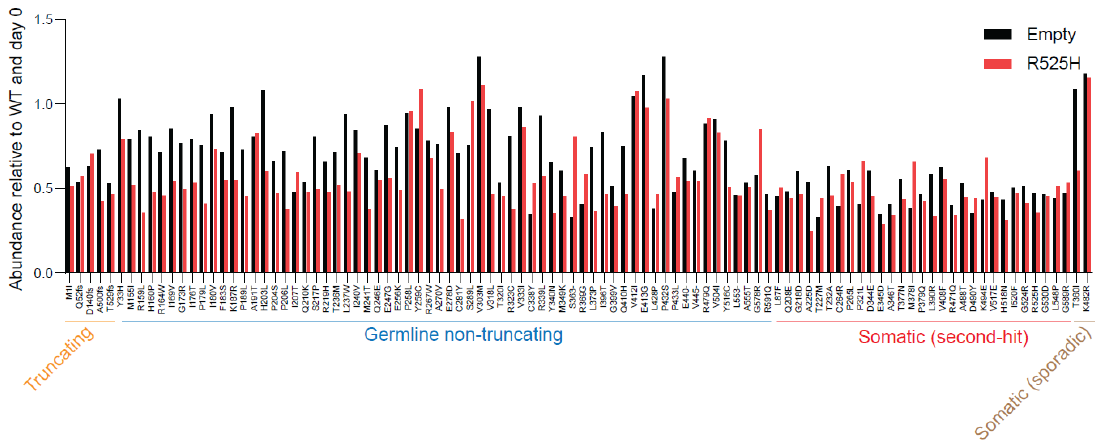

Supplementary figure 4. Abundance of germline and somatic DDX41 variants relative to day 0 and DDX41 wild-type variants in control and p.R525H cells.

Supplementary Table 1. List of variants tested in various systems and assays.

|  | **P/LP or VUS** | **Rescue**  ***Ddx41^f/f^* Lin-cells and cell line** | **MOLM13** | **C57Bl/6 Mice** | **Effect with combined with p.R525H** | **Observed with p.R525H in patients?** |
| --- | --- | --- | --- | --- | --- | --- |
| **K9E** | VUS | ✔️ | ✔️ |  |  |  |
| **Y33H** | VUS | ✔️ | ✔️ |  |  |  |
| **Q52fs** | Trunc | **X** | **X** |  |  | ✔️ |
| **D140fs** | Trunc | **X** | **X** |  |  | ✔️ |
| **M155H/I** | VUS | ✔️ | ✔️ |  | ↓ |  |
| **R164W** | P/LP | ✔️ | x |  | ↓ |  |
| **G173R** | P/LP | ✔️ | ✔️ | Het Only | ↓ | ✔️ |
| **R219H** | P/LP | ✔️ | ✔️ |  | ↓ | ✔️ |
| **P258L** | P/LP | ✔️ | x |  | ↓ | ✔️ |
| **Y259C** | P/LP | ✔️ | Slow growth | Het Only | ↓ | ✔️ |
| **K331del** | P/LP | ✔️ | x |  | ↓ | ✔️ |
| **I396T** | P/LP | ✔️ | ✔️ | ✔️ | ↓ | ✔️ |
| **R525H** | Somatic | X |  |  |  |  |
